# Co-localization of pain and reduced intra-epidermal nerve fiber density in individuals with HIV-associated sensory neuropathy

**DOI:** 10.1101/587345

**Authors:** Imraan G Patel, Peter R Kamerman

## Abstract

**Introduction:** There is poor correlation between decreases in intra-epidermal nerve fiber density (IENFD) and the presence of pain in HIV-associated sensory neuropathy (HIV-SN) and other painful distal symmetrical polyneuropathies.

**Objective:** We investigated whether in individuals with HIV-SN, having pain at the ankle skin biopsy site was associated with lower IENFD compared to when there was no pain at the ankle biopsy site.

**Methods:** We recruited 15 individuals with symptomatic HIV-SN. Nine had pain at the site where the ankle biopsy was taken, while six did not. Skin punch biopsies for IENFD quantification were taken from the ankle and the thigh. Contrasts between the two groups were made using the overlap of confidence interval (CI) method.

**Results:** IENFD was substantially lower in the group that had pain at the site of the ankle biopsy compared to the other group [6.6 (CI: 5.3 to 7.2) vs. 3.3 (CI: 10.0 to 15.0) fibers/mm]. However, there was no group differences at the thigh biopsy site [15.6 (CI: 15.0 to 15.9) vs 16.2 (CI: 14.5 to 17.8) fibers/mm]. When taking the ratio of ankle IENFD:thigh IENFD, the point estimate for the pain at the ankle group [0.43 (CI: 0.36 to 0.48)] was about half that of the other group [0.81 (CI: 0.68 to 0.87)].

**Conclusion:** Thus, co-localization of pain to the ankle is associated with meaningful decreases in ankle IENFD.

**Summary:** Having pain at the ankle biopsy site is associated with lower intra-epidermal nerve fiber density compared to not having pain at the ankle biopsy site.

## Introduction

HIV-associated sensory neuropathy (HIV-SN) is distal symmetrical polyneuropathy associated with HIV infection and its treatment [7]. Like other peripheral neuropathies, it is often, but not always painful, and it is unclear what determines whether the phenotype with pain develops [4,8,16]. There have been several efforts to correlate the presence of pain in HIV-SN to objective measures of a peripheral nerve lesion Quantitative sensory testing on the dorsum of the foot has shown that compared to non-painful HIV-SN, painful HIV-SN is associated with mechanical allodynia and mechanical hyperalgesia [1]. While intra-epidermal nerve fiber density (IENFD) is reduced in HIV-SN and is associated with pain intensity in those individuals with pain, no associations have been identified between IENFD and the presence of asymptomatic or symptomatic HIV-SN [14,17,20]. However, lower ankle IENFD was associated with increased risk of transitioning from asymptomatic to symptomatic HIV-SN [6]. We hypothesized that this tangential evidence for an association between IENFD and the presence of pain in HIV-SN results from the anatomical separation of where pain may be occuring (typically the feet) [13], and where distal skin biopsies for IENFD quantification are taken (the ankle) [11,12]. We therefore undertook an exploratory study assessing whether in individuals with HIV-SN, having pain at the ankle biopsy site was associated with lower IENFD at the ankle compared to having no pain or pain restricted to the feet.

## Methods

The study was approved by the Human Research Ethics Committee of the University of the Witwatersrand.

### Study population

We recruited a convenience sample of people living with HIV/AIDS and who were attending the Green House Pharmacy at Chris-Hani Baragwanath Hospital, Johannesburg, South Africa. Study inclusion criteria were: i) being on stable combination antiretroviral therapy (cART) for at least six months; ii) having a current CD4 T-cell count > 350 cells/mm^3^; iii) fulfilling our diagnostic criteria for the presence of distal symmetrical polyneuropathy; iv) the onset of signs or symptoms of neuropathy occurred after the initiation of cART; v) not having diabetes mellitus, and vi) not having any current skin lesions on the lower legs. Thirty-five (35) people gave informed consent and were screened. Fifteen (15) met the inclusion requirements.

### Neuropathy assessment

We assessed for symptomatic neuropathy in the lower limbs using the ACTG Brief Peripheral Neuropathy Screen (BPNS) [2], which has a case definition of the bilateral presence of at least one neurological sign (reduced or absent vibration detection in the great toes, absent ankle reflexes), and at least one symptom of polyneuropathy (pain, numbness, paresthesias) in a consistent neuroanatomical pattern. The use of medical history and the BPNS places the diagnostic certainty of our case definition at the level of probable painful neuropathy [3].

Once neuropathy had been diagnosed, the presence, severity and location of any symptoms of pain were assessed using a guided interview. Pain intensity was assessed using an 11-point numerical pain rating scale. Depending on the outcome of the assessment of pain, participants were grouped into a *‘pain’* group (n = 9) or a *‘no pain’* group (n = 6). Participants in the *‘no pain’* group met our case definition for neuropathy, but did not report experiencing pain at the ankle or thigh biopsy sites (two participants had no pain at all, but had numbness or paresthesias, and four had pain that was limited to the soles of their feet). Participants in the *‘pain group’* met our case definition for neuropathy and they had pain that extended proximally from the foot to at least the ankle biopsy site, but not past the knee.

### Skin biopsies

Three-millimeter (3mm) skin punch biopsies were taken from the ankle (10cm above the lateral malleolus) and from the lateral aspect of the proximal thigh. The thigh biopsies served as a reference site as this site is only affected very late in the progression of HIV-associated sensory neuropathy [19]. We followed published methods for the biopsy procedure, tissue fixing and preparation, panaxonal marker protein gene product (PGP) 9.5 staining, and fiber density quantification [9,11,12]. Three randomly chosen sections were assessed per biopsy, and the single assessor (IGP) was blinded to the source of the biopsy (*‘pain’* vs *‘no pain’*, and thigh vs ankle).

### Data analysis

We assessed the following: i) the proportion of ankle biopsies with fiber densities below the normal range, ii) the fiber densities at the ankle and at the thigh, and iii) the ankle:thigh fiber density ratios. Because of the small sample size, we chose to conducted exploratory analysis using the overlap of confidence intervals method. That is, failure of confidence intervals to overlap indicates a significant and meaningful difference. The confidence interval method of significance testing requires the confidence level of the intervals be adjusted to maintain a 5% error rate [5,10]. Bias-corrected accelerated confidence intervals were generating using bootstrapping, with 100000 replicates. All analyses were performed in the R Statistical Environment (R v3.5.2) [15]. A full analysis script is provided in as supplementary material (Supplement 1).

## Results

### Description of the groups

The sex ratio (female:male) was 3:3 in the *‘no pain’* group and 7:2 in the *‘pain’* group. Participants in the two groups were of similar age [*‘pain’* group (years): mean = 41, SD = 10; *‘no pain’* group: mean = 43, SD = 4; 95% confidence interval for the difference in location = −5 to 11 years], and had similar current CD4 T-cell counts [*‘pain’* group (cells/mm^3^): median = 469, range = 353 to 653; *‘no pain’* group: median = 543, range = 368 to 770; 95% confidence interval for the difference in location = −90 to 205 cells/mm^3^]. In the *‘pain’* group, median pain intensity was 10 (range: 7 to 10), and the four participants with foot pain in the *‘no pain’* group had a median pain intensity of 5 (range: 1-10).

### Intraepithelial nerve fiber density

Figure 1 shows IENFD data for the *‘pain’* group and the *‘no pain’* group at the ankle and thigh, and the ankle:thigh IENFD ratio for the groups. All participants in the *‘pain’* group (9/9) had ankle fiber densities below the 5th percentile for healthy adults, whereas only two (33%) of individuals in the *‘no pain’* group (2/6) had values below the 5th percentile (Lauria et al. 2010).

**Figure 1.**
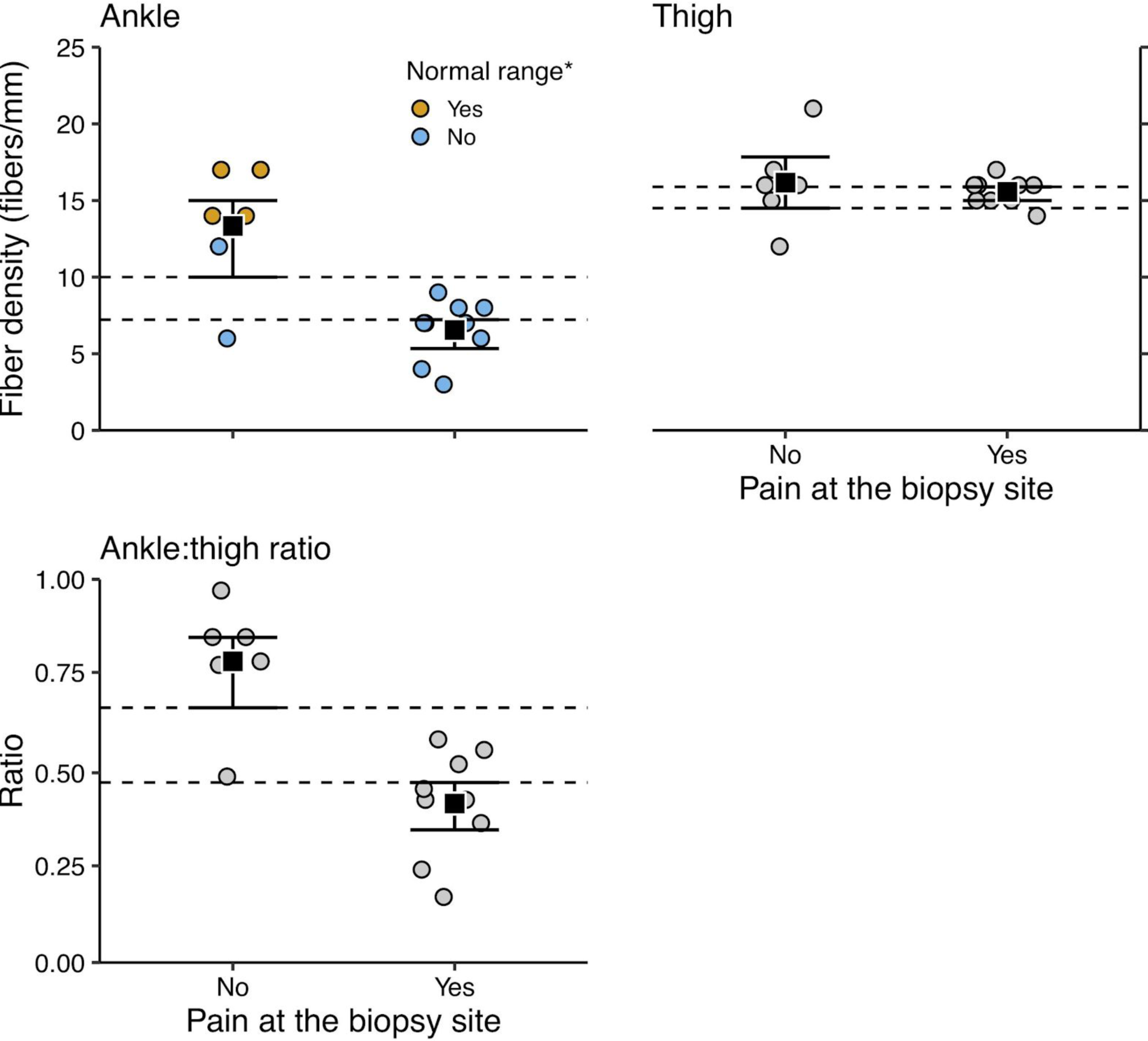
Mean and confidence interval (CI) of intraepidermal fiber densities (IENFD) for the *‘pain’* group (pain at biopsy site: yes) and the *‘no pain’* group (pain at biopsy site: no) at the ankle (top left panel) and the thigh (top right panel), and the ankle:thigh IENFD ratio (bottom left panel). Confidence intervals were calculated using bootstrapping and the confidence limits reflect 95% confidence intervals adjusted to maintain an 5% long-term error rate (see the data analysis section for details). Absence of overlap between confidence intervals indicates the the data are compatible with a statistically significant and a meaningful difference between groups. At the ankle, there was no overlap of the point estimates and confidence intervals, with the *‘pain’* group having a mean of 6.6 (85.9% CI: 5.3 to 7.2) fibers/mm, while the *‘no pain’* group had a mean of 13.3 (85.9% CI: 10.0 to 15.0) fibers/mm. So the *‘pain’* group had a point estimate of about half the fiber density of the *‘no pain’* group, and the upper limit of the confidence interval for the *‘pain’* group was about 2.5 fibers/mm different from that of the lower limit of the confidence interval of the *‘no pain’* group. In contrast, there was substantial overlap in the confidence intervals at the thigh, with the *‘pain’* group having a mean of 15.6 (88.4% CI: 15.0 to 15.9) fibers/mm, and the *‘no pain’* had a similar mean of 16.2 (88.4% CI: 14.5 to 17.8) fibers/mm. The overlap between confidence intervals, and point estimates by confidence intervals indicates that the data are compatible with no notable difference between the two groups at the reference site. When expressed as the ankle:thigh IENFD ratio, the *‘pain’* group had a mean ratio of 0.43 (83.6% CI: 0.36 to 0.48), which is about half that of the *‘no pain’* group, which had a mean ratio of 0.81 (83.6% CI: 0.68 to 0.87). Thus, the ratio data reinforce the findings from the ankle data.

At the ankle (Figure 1: Ankle) the point estimates and confidence intervals of the two groups show no overlap, such that upper confidence limit in the *‘pain’* group was about 2.5 fibers/mm lower than the lower limit of the confidence interval for the *’no pain’* group, indicating that the data are compatible with a meaningful difference in IENFD between the two groups. At the thigh, there was substantial overlap in the confidence intervals for the two groups’ IENFD at the thigh (Figure 1: Thigh). Thus, at the reference site, the data are compatible with the two groups having similar IENFD. When expressed as the ratio of ankle IENFD to thigh IENFD, the *‘pain’* group had a lower ratio compared to that of the *‘no pain’* group, with no overlap in point estimates and confidence intervals of the two groups, indicating a statistically significant and a meaningful difference (Figure 1: Ankle:thigh).

## Discussion

We conducted an exploratory study assessing whether having pain at the ankle biopsy site was associated with lower IENFD at the ankle compared to having no pain or pain restricted to the feet. We report that our data support our hypothesis that the failure to identify an association between IENFD changes and the presence of painful symptoms in HIV-SN results from the anatomical separation of where pain may be occuring, and where distal skin biopsies for IENFD quantification are taken. More broadly, our findings may, in part, explain the poor diagnostic performance of IENFD in HIV-SN; a condition that typically is painful and where screening tools typically include symptom assessment [2,13,17,18,20]. That is, ankle IENFD may best correlate with the presence of symptoms when the symptoms co-localize to the place the biopsy is taken from.

Our data are consistent with the findings of a recent meta-analysis on the association between IENFD and features of distal symmetrical polyneuropathies [8]. The meta-analysis found low levels of association between IENFD and other features of neuropathy, except for when measurements and IENFD assessment were co-localized. In this case the measures were cold detection threshold and warm detection threshold. Taken together with our data, the results of the meta-analysis have broad implications for research on distal symmetrical polyneuropathies in terms of making sure that measurements of function, symptoms, and morphology target the same anatomical area.

Our study was only an small exploratory study, which included a heterogenous *‘no pain’* group. Nevertheless, we believe that we have analysed the data conservatively, and that our findings are not spurious. Indeed, our findings are consistent with the presumed neurobiology, where morphological changes should associate with symptoms in a circumscribed neuroanatomical pattern [3]. As such, we believe that there is merit in the study, and that it provides impetus for further research on the association between IENFD and co-localization of other characteristics of distal symmetrical polyneuropathies.

## Acknowledgements

The research was funded by the Rated Researchers Programme of the National Research Foundation of South Africa, and the Medical Faculty Research Endowment Fund of the Faculty of Health Sciences, University of the Witwatersrand. The authors declare no conflicts of interest.

