## Supplement 1 for "Co-localization of pain and reduced intra-epidermal nerve fiber density in individuals with HIV-associated sensory neuropathy"

Data analysis

*Peter Kamerman*

*Last updated: 23 March 2019*

**Manuscript figure** **13**

**Session information** **17**

---

#### Import data

```
data <- read_csv('data-cleaned/data.csv')
```

---

#### Tabular summaries

##### Whole cohort

```r
# All
data %>%
    mutate_if(is.character, factor) %>%
    skim() %>%
    kable()
```

Skim summary statistics
n obs: 15
n variables: 11

Variable type: factor

| variable | missing | complete | n | n_unique | top_counts | ordered |
|---|---|---|---|---|---|---|
| below_normal | 0 | 15 | 15 | 2 | yes: 11, no: 4, NA: 0 | FALSE |
| Pain_at_biopsy_site | 0 | 15 | 15 | 2 | yes: 9, no: 6, NA: 0 | FALSE |
| Pain_location | 0 | 15 | 15 | 3 | Inc: 9, Sol: 4, No : 2, NA: 0 | FALSE |
| Sex | 0 | 15 | 15 | 2 | Fem: 10, Mal: 5, NA: 0 | FALSE |

Variable type: numeric

| variable | missing | complete | n | mean | sd | p0 | p50 | p100 |
|---|---|---|---|---|---|---|---|---|
| Age | 0 | 15 | 15 | 41.67 | 8.13 | 25 | 41 | 54 |
| ankle_thigh_ratio | 0 | 15 | 15 | 0.58 | 0.24 | 0.18 | 0.53 | 1 |
| Current_CD4 | 0 | 15 | 15 | 510.07 | 116.44 | 353 | 492 | 770 |
| IENFD_ankle | 0 | 15 | 15 | 9.27 | 4.46 | 3 | 8 | 17 |
| IENFD_thigh | 0 | 15 | 15 | 15.8 | 1.9 | 12 | 16 | 21 |
| normative_ankle | 0 | 15 | 15 | 10.87 | 1.36 | 8.9 | 11.2 | 13.5 |
| Pain_intensity | 0 | 15 | 15 | 7.07 | 3.92 | 0 | 10 | 10 |

##### By group

```r
data %>%
    mutate_if(is.character, factor) %>%
    group_by(Pain_at_biopsy_site) %>%
    skim() %>%
    kable()
```

Skim summary statistics
n obs: 15
n variables: 11

Variable type: factor

| Pain_at_biopsy_site | variable | missing | complete | n | n_unique | top_counts | ordered |
|---|---|---|---|---|---|---|---|
| no | below_normal | 0 | 6 | 6 | 2 | no: 4, yes: 2, NA: 0 | FALSE |
| no | Pain_location | 0 | 6 | 6 | 2 | Sol: 4, No : 2, Inc: 0, NA: 0 | FALSE |
| no | Sex | 0 | 6 | 6 | 2 | Fem: 3, Mal: 3, NA: 0 | FALSE |
| yes | below_normal | 0 | 9 | 9 | 1 | yes: 9, no: 0, NA: 0 | FALSE |
| yes | Pain_location | 0 | 9 | 9 | 1 | Inc: 9, No : 0, Sol: 0, NA: 0 | FALSE |
| yes | Sex | 0 | 9 | 9 | 2 | Fem: 7, Mal: 2, NA: 0 | FALSE |

Variable type: numeric

| Pain_at_biopsy_site | variable | missing | complete | n | mean | sd | p0 | p50 | p100 |
|---|---|---|---|---|---|---|---|---|---|
| no | Age | 0 | 6 | 6 | 43.33 | 4.27 | 38 | 43.5 | 49 |
| no | ankle_thigh_ratio | 0 | 6 | 6 | 0.81 | 0.17 | 0.5 | 0.84 | 1 |
| no | Current_CD4 | 0 | 6 | 6 | 551.5 | 145.05 | 368 | 543.5 | 770 |
| no | IENFD_ankle | 0 | 6 | 6 | 13.33 | 4.08 | 6 | 14 | 17 |
| no | IENFD_thigh | 0 | 6 | 6 | 16.17 | 2.93 | 12 | 16 | 21 |
| no | normative_ankle | 0 | 6 | 6 | 10.6 | 1.18 | 9.6 | 10.4 | 12.4 |
| no | Pain_intensity | 0 | 6 | 6 | 3.5 | 3.94 | 0 | 3 | 10 |
| yes | Age | 0 | 9 | 9 | 40.56 | 10.04 | 25 | 40 | 54 |
| yes | ankle_thigh_ratio | 0 | 9 | 9 | 0.43 | 0.14 | 0.18 | 0.44 | 0.6 |
| yes | Current_CD4 | 0 | 9 | 9 | 482.44 | 91.83 | 353 | 469 | 653 |
| yes | IENFD_ankle | 0 | 9 | 9 | 6.56 | 1.94 | 3 | 7 | 9 |
| yes | IENFD_thigh | 0 | 9 | 9 | 15.56 | 0.88 | 14 | 16 | 17 |
| yes | normative_ankle | 0 | 9 | 9 | 11.06 | 1.5 | 8.9 | 11.2 | 13.5 |
| yes | Pain_intensity | 0 | 9 | 9 | 9.44 | 1.13 | 7 | 10 | 10 |

##### No pain group: painful feet vs no pain at all

```
data %>%
    filter(Pain_at_biopsy_site == 'no') %>%
    group_by(Pain_location) %>%
    skim() %>%
    kable()
```

Skim summary statistics
n obs: 6
n variables: 11

Variable type: character

| Pain_location | variable | missing | complete | n | min | max | empty | n_unique |
|---|---|---|---|---|---|---|---|---|
| No pain | below_normal | 0 | 2 | 2 | 2 | 2 | 0 | 1 |
| No pain | Pain_at_biopsy_site | 0 | 2 | 2 | 2 | 2 | 0 | 1 |
| No pain | Sex | 0 | 2 | 2 | 6 | 6 | 0 | 1 |
| Sole of feet only | below_normal | 0 | 4 | 4 | 2 | 3 | 0 | 2 |
| Sole of feet only | Pain_at_biopsy_site | 0 | 4 | 4 | 2 | 2 | 0 | 1 |

| Pain_location | variable | missing | complete | n | min | max | empty | n_unique |
|---|---|---|---|---|---|---|---|---|
| Sole of feet only | Sex | 0 | 4 | 4 | 4 | 6 | 0 | 2 |

Variable type: numeric

| Pain_location | variable | missing | complete | n | mean | sd | p0 | p50 | p100 |
|---|---|---|---|---|---|---|---|---|---|
| No pain | Age | 0 | 2 | 2 | 43 | 4.24 | 40 | 43 | 46 |
| No pain | ankle_thigh_ratio | 0 | 2 | 2 | 0.84 | 0.046 | 0.81 | 0.84 | 0.88 |
| No pain | Current_CD4 | 0 | 2 | 2 | 652.5 | 166.17 | 535 | 652.5 | 770 |
| No pain | IENFD_ankle | 0 | 2 | 2 | 15.5 | 2.12 | 14 | 15.5 | 17 |
| No pain | IENFD_thigh | 0 | 2 | 2 | 18.5 | 3.54 | 16 | 18.5 | 21 |
| No pain | normative_ankle | 0 | 2 | 2 | 11.2 | 0 | 11.2 | 11.2 | 11.2 |
| No pain | Pain_intensity | 0 | 2 | 2 | 0 | 0 | 0 | 0 | 0 |
| Sole of feet only | Age | 0 | 4 | 4 | 43.5 | 4.93 | 38 | 43.5 | 49 |
| Sole of feet only | ankle_thigh_ratio | 0 | 4 | 4 | 0.79 | 0.21 | 0.5 | 0.84 | 1 |
| Sole of feet only | Current_CD4 | 0 | 4 | 4 | 501 | 125.14 | 368 | 493 | 650 |
| Sole of feet only | IENFD_ankle | 0 | 4 | 4 | 12.25 | 4.65 | 6 | 13 | 17 |
| Sole of feet only | IENFD_thigh | 0 | 4 | 4 | 15 | 2.16 | 12 | 15.5 | 17 |
| Sole of feet only | normative_ankle | 0 | 4 | 4 | 10.3 | 1.4 | 9.6 | 9.6 | 12.4 |
| Sole of feet only | Pain_intensity | 0 | 4 | 4 | 5.25 | 3.69 | 1 | 5 | 10 |

---

#### 95% CI for the difference in location

##### Age

```
with(data, t.test(Age ~ Pain_at_biopsy_site))
```

```
##
##  Welch Two Sample t-test
##
## data:  Age by Pain_at_biopsy_site
## t = 0.73606, df = 11.573, p-value = 0.4764
## alternative hypothesis: true difference in means is not equal to 0
## 95 percent confidence interval:
##  -5.478537 11.034092
## sample estimates:
##  mean in group no mean in group yes
##          43.33333          40.55556
```

### CD4

```
with(data, wilcox.test(Current_CD4 ~ Pain_at_biopsy_site,
                       conf.int = TRUE))
```

```
##
##  Wilcoxon rank sum test
##
## data:  Current_CD4 by Pain_at_biopsy_site
## W = 34, p-value = 0.4559
## alternative hypothesis: true location shift is not equal to 0
## 95 percent confidence interval:
##  -90 205
## sample estimates:
## difference in location
##                   71.5
```

---

### Exploratory plots

#### Proportion within normal range

```
data %>%
    ggplot(data = .) +
    aes(Pain_at_biopsy_site,
        fill = below_normal) +
    geom_bar(position = position_fill()) +
    geom_text(stat = 'count',
              aes(y = ..prop.., label = paste0('(n = ', ..count.., ')')),
              position = position_fill(vjust = 0.5),
              size = 6) +
    labs(x = 'Pain at the biopsy site',
         y = 'Proportion',
         caption = '* Below the 50th percentile for age and sex') +
    scale_y_continuous(expand = c(0, 0)) +
    scale_x_discrete(labels = c('No', 'Yes')) +
    scale_fill_manual(values = cb_pal,
                      name = 'Normal range*',
                      labels = c('Yes', 'No')) +
    theme_bw(base_size = 16) +
    theme(panel.border = element_blank(),
          axis.line = element_line(size = 0.5),
          panel.grid = element_blank())
```

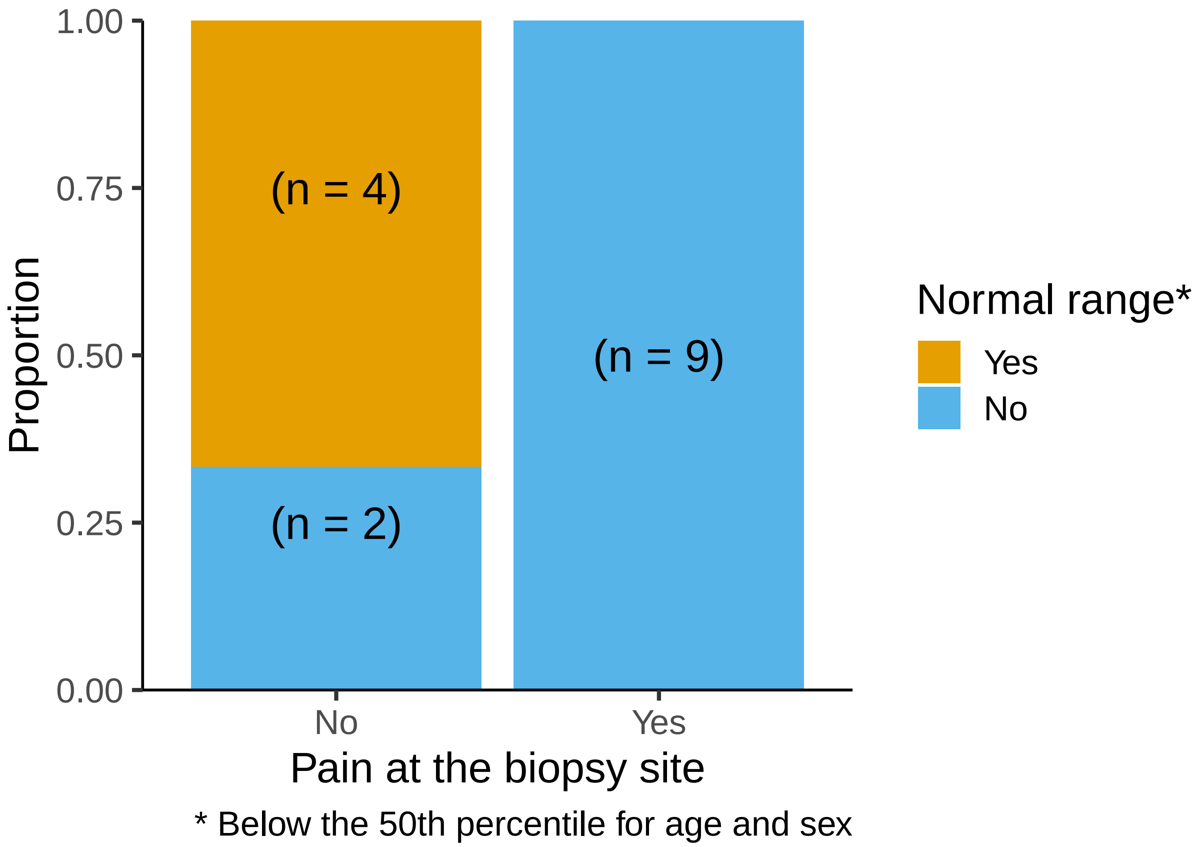

###### Summary ankle

```
data %>%
    ggplot(data = .) +
    aes(x = Pain_at_biopsy_site,
        y = IENFD_ankle) +
    geom_boxplot(fill = '#CCCCCC') +
    geom_point(aes(fill = below_normal),
               position = position_jitter(width = 0.3, height = 0),
               shape = 21,
               size = 4) +
    labs(title = 'Ankle',
         x = 'Pain at the biopsy site',
         y = 'Fiber density (fibers/mm)',
         caption = '* Below the 50th percentile for age and sex') +
    scale_y_continuous(limits = c(0, max(data$IENFD_ankle))) +
    scale_x_discrete(labels = c('No', 'Yes')) +
    scale_fill_manual(values = cb_pal,
                      name = 'Normal range*',
                      labels = c('Yes', 'No')) +
    theme_bw(base_size = 16) +
    theme(panel.border = element_blank(),
          axis.line = element_line(size = 0.5),
          panel.grid = element_blank(),
          legend.background = element_rect(colour = '#000000', size = 0.5),
```

```
      legend.position = c(0.85, 0.85))
```

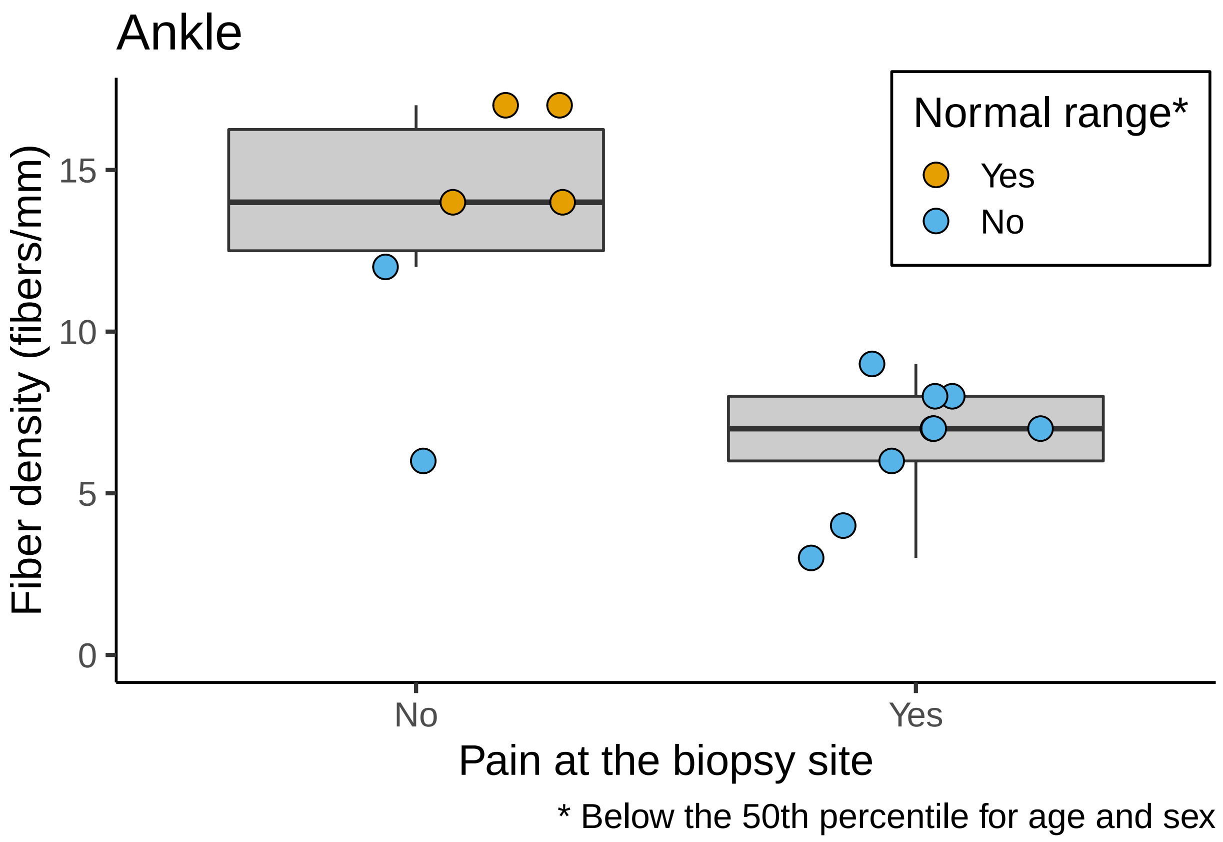

###### Summary thigh

```
data %>%
    ggplot(data = .) +
    aes(x = Pain_at_biopsy_site,
        y = IENFD_thigh) +
    geom_boxplot(fill = '#CCCCCC') +
    geom_point(fill = '#FFFFFF',
               position = position_jitter(width = 0.3, height = 0),
               shape = 21,
               size = 4,
               stroke = 0.8) +
    labs(title = 'Thigh',
         x = 'Pain at the biopsy site',
         y = 'Fiber density (fibers/mm)') +
    scale_y_continuous(limits = c(0, max(data$IENFD_thigh))) +
    scale_x_discrete(labels = c('No', 'Yes')) +
    theme_bw(base_size = 16) +
    theme(panel.border = element_blank(),
          axis.line = element_line(size = 0.5),
          panel.grid = element_blank())
```

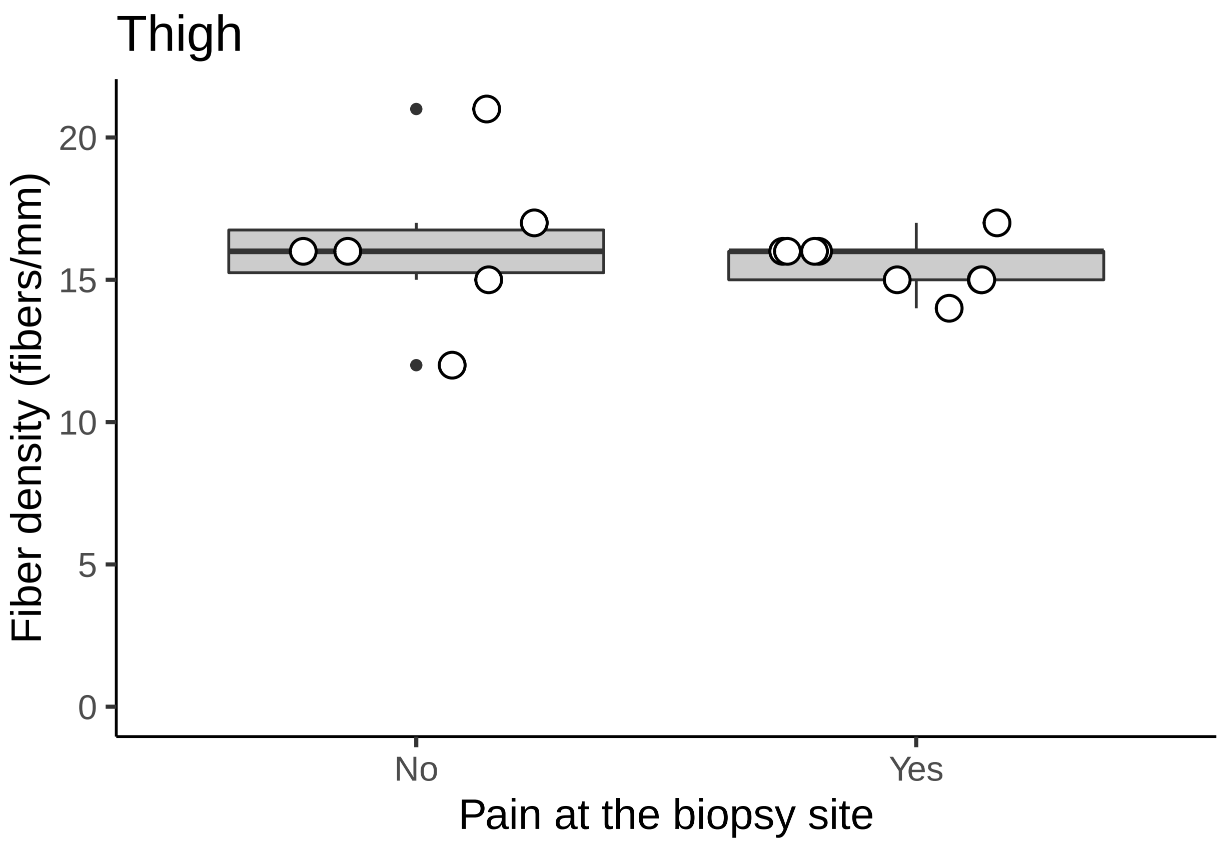

**Summary ankle:thigh ratio**

```
data %>%
    ggplot(data = .) +
    aes(x = Pain_at_biopsy_site,
        y = ankle_thigh_ratio) +
    geom_boxplot(fill = '#CCCCCC') +
    geom_point(fill = '#FFFFFF',
               position = position_jitter(width = 0.3, height = 0),
               shape = 21,
               size = 4,
               stroke = 0.8) +
    labs(title = 'Ankle:Thigh ratio',
         x = 'Pain at the biopsy site',
         y = 'Ratio') +
    scale_y_continuous(limits = c(0, max(data$ankle_thigh_ratio))) +
    scale_x_discrete(labels = c('No', 'Yes')) +
    theme_bw(base_size = 16) +
    theme(panel.border = element_blank(),
          axis.line = element_line(size = 0.5),
          panel.grid = element_blank())
```

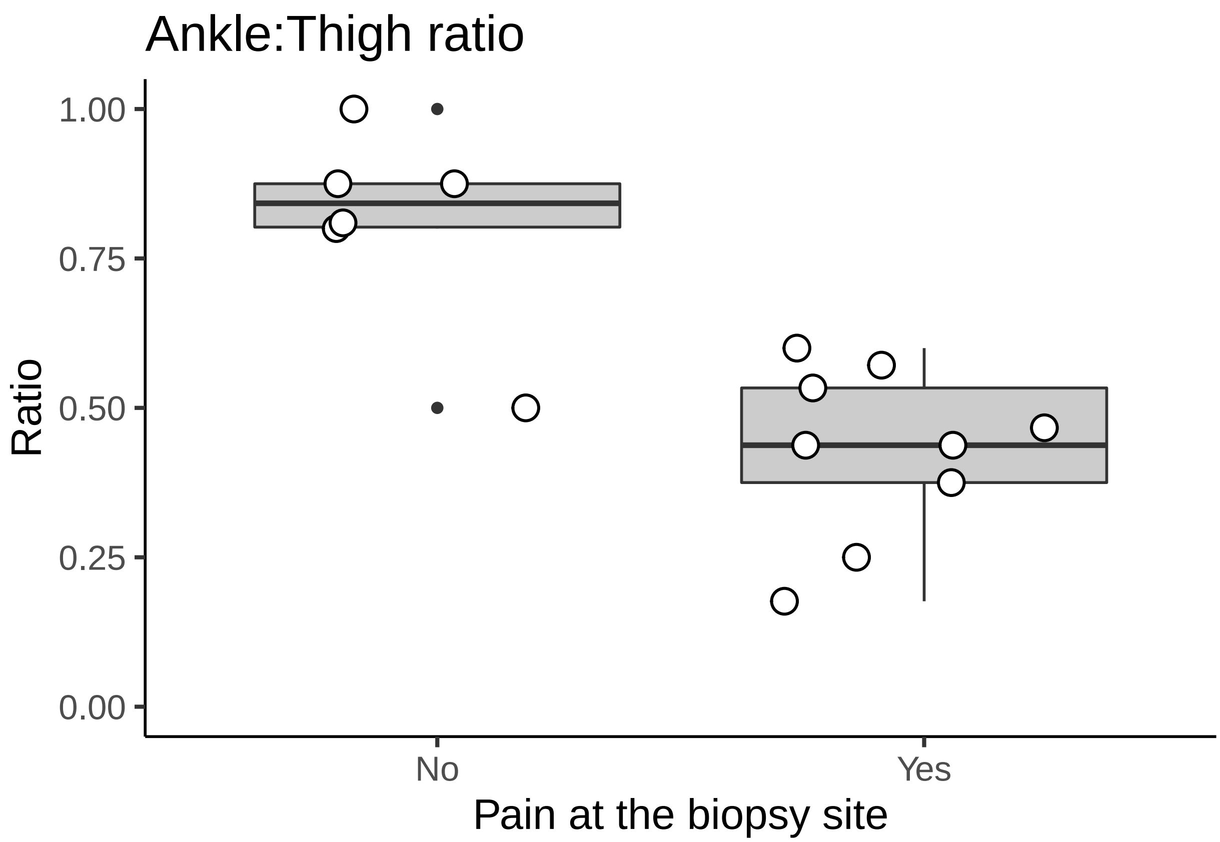

---

#### Calculate bootstrap CI for IENFDs

##### Define boot function

```
boot_mean <- function(d, i) {
    mean(d[i])
}
```

##### Ankle

```
# Extract ankle data
ankle_no <- data$IENFD_ankle[data$Pain_at_biopsy_site == 'no']
ankle_yes <- data$IENFD_ankle[data$Pain_at_biopsy_site == 'yes']

# Calculate CI width to control the type I error rate at 5%
# (Knol et al., 2011, DOI: 10.1007/s10654-011-9563-8)
p <- sd(ankle_yes)/sd(ankle_no)
z <- 1.96 * (sqrt(1 + p^2)/(1 + p))
ci <- round((1 - (2 * pnorm(-z))) * 100, 2); ci
```

```
## [1] 85.86
```

```r
# Bootstrap values
## Ankle: no pain group
set.seed(2019)
ankle_no_boot <- boot(data = ankle_no,
                      statistic = boot_mean,
                      R = 100000,
                      stype = 'i')

ankle_no_ci <- boot.ci(ankle_no_boot,
                       conf = 0.8586,
                       type = 'bca')

## Ankle: pain group
set.seed(2019)
ankle_yes_boot <- boot(data = ankle_yes,
                      statistic = boot_mean,
                      R = 100000,
                      stype = 'i')

ankle_yes_ci <- boot.ci(ankle_yes_boot,
                       conf = 0.8586,
                       type = 'bca')

# Make a dataframe
ankle <- tibble(Pain_at_biopsy_site = c('no', 'yes'),
                IENFD_mean = c(ankle_no_boot$t0, ankle_yes_boot$t0),
                IENFD_lowerCI = c(ankle_no_ci$bca[[4]], ankle_yes_ci$bca[[4]]),
                IENFD_upperCI = c(ankle_no_ci$bca[[5]], ankle_yes_ci$bca[[5]]))

# View tabulated data
ankle
```

```
## # A tibble: 2 x 4
##   Pain_at_biopsy_site IENFD_mean IENFD_lowerCI IENFD_upperCI
##   <chr>                    <dbl>         <dbl>         <dbl>
## 1 no                       13.3          10            15
## 2 yes                       6.56          5.33          7.22
```

##### Thigh

```r
# Extract thigh data
thigh_no <- data$IENFD_thigh[data$Pain_at_biopsy_site == 'no']
thigh_yes <- data$IENFD_thigh[data$Pain_at_biopsy_site == 'yes']

# Calculate CI width to control the type I error rate at 5%
# (Knol et al., 2011, DOI: 10.1007/s10654-011-9563-8)
p <- sd(thigh_yes)/sd(thigh_no)
z <- 1.96 * (sqrt(1 + p^2)/(1 + p))
```

```r
ci <- round((1 - (2 * pnorm(-z))) * 100, 2); ci
```

```
## [1] 88.43
```

```r
# Bootstrap values
## Thigh: no pain group
set.seed(2019)
thigh_no_boot <- boot(data = thigh_no,
                      statistic = boot_mean,
                      R = 100000,
                      stype = 'i')

thigh_no_ci <- boot.ci(thigh_no_boot,
                       conf = 0.8843,
                       type = 'bca')

## Thigh: pain group
set.seed(2019)
thigh_yes_boot <- boot(data = thigh_yes,
                       statistic = boot_mean,
                       R = 100000,
                       stype = 'i'); thigh_yes_boot
```

```
##
## ORDINARY NONPARAMETRIC BOOTSTRAP
##
##
## Call:
## boot(data = thigh_yes, statistic = boot_mean, R = 1e+05, stype = "i")
##
##
## Bootstrap Statistics :
##     original        bias    std. error
## t1* 15.55556 -0.0006033333   0.2776236
```

```r
thigh_yes_ci <- boot.ci(thigh_yes_boot,
                        conf = 0.8843,
                        type = 'bca'); thigh_yes_ci
```

```
## BOOTSTRAP CONFIDENCE INTERVAL CALCULATIONS
## Based on 100000 bootstrap replicates
##
## CALL :
## boot.ci(boot.out = thigh_yes_boot, conf = 0.8843, type = "bca")
##
## Intervals :
## Level       BCa
## 88.43%   (15.00, 15.89 )
## Calculations and Intervals on Original Scale
```

```r
# Make a dataframe
thigh <- tibble(Pain_at_biopsy_site = c('no', 'yes'),
```

```r
                 IENFD_mean = c(thigh_no_boot$t0, thigh_yes_boot$t0),
                 IENFD_lowerCI = c(thigh_no_ci$bca[[4]], thigh_yes_ci$bca[[4]]),
                 IENFD_upperCI = c(thigh_no_ci$bca[[5]], thigh_yes_ci$bca[[5]]))

# View tabulated data
thigh
```

```
## # A tibble: 2 x 4
##   Pain_at_biopsy_site IENFD_mean IENFD_lowerCI IENFD_upperCI
##   <chr>                    <dbl>         <dbl>         <dbl>
## 1 no                        16.2          14.5          17.8
## 2 yes                       15.6          15            15.9
```

##### Ankle:Thigh ratio

```r
# Extract ratio data
ratio_no <- data$ankle_thigh_ratio[data$Pain_at_biopsy_site == 'no']
ratio_yes <- data$ankle_thigh_ratio[data$Pain_at_biopsy_site == 'yes']

# Calculate CI width to control the type I error rate at 5%
# (Knol et al., 2011, DOI: 10.1007/s10654-011-9563-8)
p <- sd(ratio_yes)/sd(ratio_no)
z <- 1.96 * (sqrt(1 + p^2)/(1 + p))
ci <- round((1 - (2 * pnorm(-z))) * 100, 2); ci
```

```
## [1] 83.57
```

```r
# Bootstrap values
## ratio: no pain group
set.seed(2019)
ratio_no_boot <- boot(data = ratio_no,
                      statistic = boot_mean,
                      R = 100000,
                      stype = 'i')

ratio_no_ci <- boot.ci(ratio_no_boot,
                       conf = 0.8357,
                       type = 'bca')

## ratio: pain group
set.seed(2019)
ratio_yes_boot <- boot(data = ratio_yes,
                       statistic = boot_mean,
                       R = 100000,
                       stype = 'i'); ratio_yes_boot
```

```
##
## ORDINARY NONPARAMETRIC BOOTSTRAP
##
##
```

```
## Call:
## boot(data = ratio_yes, statistic = boot_mean, R = 1e+05, stype = "i")
##
##
## Bootstrap Statistics :
##      original        bias    std. error
## t1* 0.4275444 -0.0001387104  0.04460309
```

```
ratio_yes_ci <- boot.ci(ratio_yes_boot,
                        conf = 0.8357,
                        type = 'bca'); ratio_yes_ci
```

```
## BOOTSTRAP CONFIDENCE INTERVAL CALCULATIONS
## Based on 100000 bootstrap replicates
##
## CALL :
## boot.ci(boot.out = ratio_yes_boot, conf = 0.8357, type = "bca")
##
## Intervals :
## Level       BCa
## 83.57%   ( 0.3566,  0.4840 )
## Calculations and Intervals on Original Scale
```

```
# Make a dataframe
ratio <- tibble(Pain_at_biopsy_site = c('no', 'yes'),
                IENFD_mean = c(ratio_no_boot$t0, ratio_yes_boot$t0),
                IENFD_lowerCI = c(ratio_no_ci$bca[[4]], ratio_yes_ci$bca[[4]]),
                IENFD_upperCI = c(ratio_no_ci$bca[[5]], ratio_yes_ci$bca[[5]]))

# View tabulated data
ratio
```

```
## # A tibble: 2 x 4
##   Pain_at_biopsy_site IENFD_mean IENFD_lowerCI IENFD_upperCI
##   <chr>                    <dbl>         <dbl>         <dbl>
## 1 no                       0.810         0.685         0.874
## 2 yes                      0.428         0.357         0.484
```

---

### Manuscript figure

```
# Plot the ankle data
gg_ankle <- ggplot() +
    aes(x = Pain_at_biopsy_site) +
    geom_hline(yintercept = ankle$IENFD_lowerCI[[1]],
               linetype = 2,
               size = 0.8) +
    geom_hline(yintercept = ankle$IENFD_upperCI[[2]],
               linetype = 2,
```

```
            size = 0.8) +
    geom_point(data = data,
               aes(y = IENFD_ankle,
                   fill = below_normal),
               position = position_jitter(width = 0.15, height = 0),
               shape = 21,
               stroke = 1,
               size = 4) +
    geom_errorbar(data = ankle,
                  aes(ymin = IENFD_lowerCI,
                      ymax = IENFD_upperCI),
                  width = 0.4,
                  size = 1) +
    geom_point(data = ankle,
               aes(y = IENFD_mean),
               shape = 22,
               colour = '#FFFFFF',
               stroke = 1,
               fill = '#000000',
               size = 6.5) +
    labs(subtitle = 'Ankle',
         x = 'Pain at the biopsy site',
         y = 'Fiber density (fibers/mm)') +
    scale_y_continuous(limits = c(0, 25),
                       expand = c(0, 0)) +
    scale_x_discrete(labels = c('No', 'Yes')) +
    scale_fill_manual(values = cb_pal,
                      name = 'Normal range*',
                      labels = c('Yes', 'No')) +
    theme_bw(base_size = 20) +
    theme(panel.border = element_blank(),
          axis.line = element_line(size = 0.8),
          axis.text.y = element_text(colour = '#000000'),
          axis.text.x = element_blank(),
          axis.title.x = element_blank(),
          panel.grid = element_blank(),
          legend.text = element_text(size = 14),
          legend.title = element_text(size = 16),
          legend.position = c(0.8, 0.85))

# Plot the thigh data
gg_thigh <- ggplot() +
    aes(x = Pain_at_biopsy_site) +
    geom_hline(yintercept = thigh$IENFD_lowerCI[[1]],
               linetype = 2,
               size = 0.8) +
    geom_hline(yintercept = thigh$IENFD_upperCI[[2]],
               linetype = 2,
               size = 0.8) +
```

```
    geom_point(data = data,
               aes(y = IENFD_thigh),
               position = position_jitter(width = 0.15,
                                          height = 0),
               shape = 21,
               size = 4,
               stroke = 1,
               fill = '#CCCCCC') +
    geom_errorbar(data = thigh,
                  aes(ymin = IENFD_lowerCI,
                      ymax = IENFD_upperCI),
                  width = 0.4,
                  size = 1) +
    geom_point(data = thigh,
               aes(y = IENFD_mean),
               shape = 22,
               colour = '#FFFFFF',
               stroke = 1,
               fill = '#000000',
               size = 6.5) +
    labs(subtitle = 'Thigh',
         x = 'Pain at the biopsy site',
         y = 'Fiber density (fibers/mm)') +
    scale_y_continuous(position = 'right',
                       limits = c(0, 25),
                       expand = c(0, 0)) +
    scale_x_discrete(labels = c('No', 'Yes')) +
    theme_bw(base_size = 20) +
    theme(panel.border = element_blank(),
          axis.line = element_line(size = 0.8),
          axis.text.x = element_text(colour = '#000000'),
          axis.text.y = element_blank(),
          axis.title.y = element_blank(),
          panel.grid = element_blank())

# Plot the ratio data
gg_ratio <- ggplot() +
    aes(x = Pain_at_biopsy_site) +
    geom_hline(yintercept = ratio$IENFD_lowerCI[[1]],
               linetype = 2,
               size = 0.8) +
    geom_hline(yintercept = ratio$IENFD_upperCI[[2]],
               linetype = 2,
               size = 0.8) +
    geom_point(data = data,
               aes(y = ankle_thigh_ratio),
               position = position_jitter(width = 0.15,
                                          height = 0),
               shape = 21,
```

```
               size = 4,
               stroke = 1,
               fill = '#CCCCCC') +
    geom_errorbar(data = ratio,
                  aes(ymin = IENFD_lowerCI,
                      ymax = IENFD_upperCI),
                  width = 0.4,
                  size = 1) +
    geom_point(data = ratio,
               aes(y = IENFD_mean),
               shape = 22,
               colour = '#FFFFFF',
               stroke = 1,
               fill = '#000000',
               size = 6.5) +
    labs(subtitle = 'Ankle:thigh ratio',
         x = 'Pain at the biopsy site',
         y = 'Ratio') +
    scale_y_continuous(position = 'left',
                       limits = c(0, 1.03),
                       labels = c('0.00', '0.25', '0.50', '0.75', '1.00'),
                       breaks = c(0, 0.26, 0.51, 0.78, 1.03),
                       expand = c(0, 0)) +
    scale_x_discrete(labels = c('No', 'Yes')) +
    theme_bw(base_size = 20) +
    theme(panel.border = element_blank(),
          axis.line = element_line(size = 0.8),
          axis.text = element_text(colour = '#000000'),
          panel.grid = element_blank())

#- Make and save the figure ---#
## Patch components together
gg_top <- gg_ankle + gg_thigh
gg_bottom <- gg_ratio
gg_patch <- gg_top + gg_bottom + plot_layout(nrow = 2)

## Save the figure
if(!dir.exists('outputs')) {
    dir.create('outputs')
    ggsave(filename = 'outputs/figure_1.pdf',
           plot = gg_patch,
           width = 10,
           height = 9)
    ggsave(filename = 'outputs/figure_1.png',
           plot = gg_patch,
           width = 10,
           height = 9)
} else {
    ggsave(filename = 'outputs/figure_1.pdf',
```

```r
        plot = gg_patch,
        width = 10,
        height = 9)
  ggsave(filename = 'outputs/figure_1.png',
        plot = gg_patch,
        width = 10,
        height = 9)
}
```

---

#### Session information

```r
sessionInfo()
```

```
## R version 3.5.2 (2018-12-20)
## Platform: x86_64-apple-darwin15.6.0 (64-bit)
## Running under: macOS Mojave 10.14.3
##
## Matrix products: default
## BLAS: /Library/Frameworks/R.framework/Versions/3.5/Resources/lib/libRblas.0.dylib
## LAPACK: /Library/Frameworks/R.framework/Versions/3.5/Resources/lib/libRlapack.dylib
##
## locale:
## [1] en_US.UTF-8/en_US.UTF-8/en_US.UTF-8/C/en_US.UTF-8/en_US.UTF-8
##
## attached base packages:
## [1] stats     graphics  grDevices utils     datasets  methods   base
##
## other attached packages:
##  [1] boot_1.3-20     skimr_1.0.5     patchwork_0.0.1 forcats_0.4.0
##  [5] stringr_1.4.0   dplyr_0.8.0.1   purrr_0.3.1     readr_1.3.1
##  [9] tidyr_0.8.3     tibble_2.0.1    ggplot2_3.1.0   tidyverse_1.2.1
##
## loaded via a namespace (and not attached):
##  [1] tidyselect_0.2.5 xfun_0.5         haven_2.1.0      lattice_0.20-38
##  [5] colorspace_1.4-0 generics_0.0.2   htmltools_0.3.6  yaml_2.2.0
##  [9] utf8_1.1.4       rlang_0.3.1      pillar_1.3.1     glue_1.3.0
## [13] withr_2.1.2.9000 modelr_0.1.4     readxl_1.3.0     plyr_1.8.4
## [17] munsell_0.5.0    gtable_0.2.0     cellranger_1.1.0 rvest_0.3.2
## [21] evaluate_0.13    labeling_0.3     knitr_1.21       fansi_0.4.0
## [25] highr_0.7        broom_0.5.1      Rcpp_1.0.0       scales_1.0.0
## [29] backports_1.1.3  jsonlite_1.6     hms_0.4.2        digest_0.6.18
## [33] stringi_1.3.1    grid_3.5.2       cli_1.0.1        tools_3.5.2
## [37] magrittr_1.5     lazyeval_0.2.1   crayon_1.3.4     pkgconfig_2.0.2
## [41] xml2_1.2.0       lubridate_1.7.4  assertthat_0.2.0 rmarkdown_1.11
## [45] httr_1.4.0       rstudioapi_0.9.0 R6_2.4.0         nlme_3.1-137
## [49] compiler_3.5.2
```
